# A Hydrodynamic Approach To Cancer

**DOI:** 10.1101/014696

**Authors:** Alexander Pinkowski, Walter Lilienblum

## Abstract

This is the pre-print version of a paper submitted to Technische Mechanik (ISSN 0232-3869)

## Introduction

In this paper we analyze several aspects of blood rheology as they relate to cancer research, with special emphasis on the hydrodynamic impact of drag-reducing agents (DRA), i.e., drag-reducing polymers (DRP) and drag-reducing surfactants (DRS).

Blood is the most important body fluid. Its rheology is non-Newtonian, meaning that blood has a non-linear shear-stress to shear-rate relationship, i.e., one has to apply a threshold force, the “yield stress”, before it moves at all. This property is determined by the composition of blood, and by the particular qualities of its components.

Blood consists predominately of plasma, with its near-Newtonian flow properties (except in elongational flow (Brust, 2013), and red blood cells (RBCs). This leads to a two-phased flow behavior, with the plasma acting as a carrier phase and the RBCs as a liquid-droplet-like carried phase suspended therein. At low shear rates, i.e., low velocity gradients, RBCs tend to form rouleaux structures which are potential sources of vortices. Low shear rates may also cause the primary, randomly scattered rouleaux to group together to create secondary rouleaux formations (cf. fig. 1, Kulicke, 1986).

**Fig. 1:**
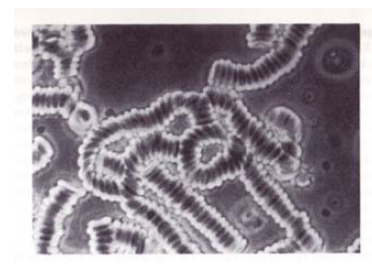
Secondary RBC rouleaux (Kulicke, 1986)

Low shear rates occur also in tumor vessels and may facilitate rouleaux formation even at low hematocrits (Sevick et al., 1989).

Blood flow is laminar below Reynolds numbers Re ≈ 2300 and turbulent for Re > 3000 (1)

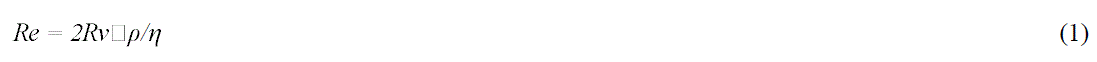

where *R* = vessel radius; *v*□ = mean velocity of blood flow (mean in time and cross-section); *ρ* = density of whole blood, and *η* = dynamic viscosity.

Hence turbulent flow tends to occur where *R* is large (in the aorta, for example), or *v*□ is large, such as during high cardiac output. However, turbulent flow may also occur where the local vessel diameter is reduced (such as in an arterial stenosis), resulting in a localized increase in blood flow velocity. Low blood viscosity *η*, such as occurs in anemia (low RBC content), predisposes to turbulence. Vessel branching (bifurcation) and vessel bending with plasma skimming (Boron et al., 2005) can also be a cause of local turbulence. In turbulent flow the parabolic velocity profile becomes blunted.

It is well known that atherosclerotic lesions of vessels are not randomly distributed but tend to occur at sites where the laminar blood flow is disturbed, such as at vessel bifurcations. Even in a healthy circulatory system this will inevitably cause endothelial lesions due to flow-induced apoptosis. Such lesions may represent potential docking points for metastasizing tumor cells.

In addition to the effects of branching and curving, vessel dilatation above critical Reynolds numbers may cause pockets of blood pooling in the low shear environment near the internal wall of blood vessels. Thus a vicious cycle can develop where increased viscosity leads to decreased flow, which in turn increases viscosity and decreases flow further, culminating in stasis and thrombosis.

Blood viscosity is measured by means of viscometers also called rheometers when examining liquids whose viscosities vary with flow conditions. So measured, blood viscosity (*η*) is always an average, effective, over-all, or apparent viscosity. Its value depends on fibrinogen concentration, hematocrit, vessel radius, temperature, and velocity profile in a narrow gap. At low flow velocities whole blood exhibits a non-Newtonian viscous-elastic behavior caused mainly by the interaction of fibrinogen with RBCs.

RBCs tend to accumulate in the vessel center, leaving RBC-depleted plasma in the wall region and thereby reducing blood viscosity in the area near the vessel wall. As a result, the near-wall velocity profile becomes steeper while it becomes smaller at the vessel center. Fibrinogen-aided RBC rouleaux formation occurring at low shear rates in the vessel center disappears at higher shear rates. While almost negligible in healthy circulatory systems, the risk of cluster formation is especially high in cases of impeded circulation since the cluster formation time is around 10 s, or about 10 times higher than heart pulse time (Oertel, 2008).

<u>Two-phase blood flow:</u> Blood can be viewed as either an emulsion or a suspension. As an emulsion, the plasma is the fluid that carries liquid RBC droplets. Seen as a suspension, RBCs are solids carried within the liquid plasma. Both emulsions and suspensions need stabilizers - emulsifiers and surfactants respectively. Surfactants serve primarily as wetting agents by reducing the surface tension of the liquid carrier surrounding the suspended solid particle. It should be mentioned that surfactant micelles orient themselves in the direction of flow. With emulsifiers (or emulgents), the following three mechanisms of action are discussed:

Surface tension theory: according to this theory emulsification takes place through the reduction of interfacial tension between two phases.

Repulsion theory: the emulsifying agent creates a film over one phase that forms globules, which repel each other. This repulsive force causes them to remain suspended in the dispersion medium.

Viscosity modification: emulgents such as polyethylene oxide (PEO) increase the viscosity of the medium, which helps create and maintain the suspension of RBC globules as dispersed phase.

The stabilizing mechanisms of surfactant and DRP could be different. With surfactants RBC aggregation is probably avoided due to the better wettability of RBCs, while DRP prevent plasma skimming by increasing local viscosity around RBCs.

It was stated (Dintenfass, 1985) that at high shear rates blood behaves like an emulsion, in which blood flow is determined by the differences in viscosity between plasma and RBCs. At low shear rates, however, blood behaves like a suspension of rigid RBCs in the carrier phase plasma. As previously mentioned, possible RBC aggregation at low shear rates is mainly of interest in affected blood circulation (Oertel, l.c.).

<u>Angiogenesis and cancer:</u> After a circulating tumor cell docks on a site in the circulatory system characterized by a favorable nutrient supply, the tumor cell penetrates the epithelial vessel layer in order to grow outside the vessel and form a larger tumor. Epithelial layers contain no blood vessels, so they must receive nourishment via diffusion of substances from the underlying connective tissue, passing through the basement membrane. The basement membrane acts as a mechanical barrier, preventing malignant cells from invading the deeper tissues.

Tumor cells can penetrate blood or lymphatic vessels, circulate through the intravascular stream, and then proliferate at another site, i.e. metastasize. For the metastatic spread of cancer tissue, growth of the vascular network is important. The formation of new blood and lymphatic vessels, i.e., angiogenesis and lymphangiogenesis, plays an essential role in the formation of a new vascular network to supply nutrients, oxygen and immune cells, and remove waste products. New growth in the vascular network is critical to the proliferation and metastatic spread of cancer cells whose growth is dependent upon an abundant supply of nutrients. Angiogenesis is regulated by both activator and inhibitor molecules. Unlike normal blood vessels, tumor blood vessels are dilated with an irregular shape.

<u>Cancer and blood rheology:</u> Angiogenesis not only supplies the nutrients needed for tumor growth, but the new vessels created in the process also serve as gateways for detached tumor cells to enter into circulation. But angiogenesis may also play a third role: new vessels are inevitably sources of micro-turbulence, creating areas with enhanced mass transfer, i.e., nutrient supply. While DRA will neither hinder angiogenesis nor prevent the escape of single tumor cells, their injection may well dampen local flow disturbances, thereby diminishing the extra nutrient supply and hindering the settlement of circulating cancer cells, i.e., metastasis.

That blood rheology in patients with ovarian cancer is impaired due to inflammatory processes is well known. More recently it was shown that the disaggregation threshold of RBCs is enhanced in woman suffering from ovarian cancer (El Bouhmadi et al. 2000). It was also demonstrated (von Tempelhoff, 2000; von Tempelhoff, Niemann et al., 1998; von Tempelhoff, Heilmann et al., 1998) that in patients with gynecologic malignancies an increase in coagulation activation occurs. With impaired blood rheology microcirculatory blood flow is reduced, in turn increasing the probability of thrombosis and promoting tumor progression and metastasis. Furthermore, in ovarian cancer patients who developed deep vein thrombosis postoperatively and during chemotherapy, the thrombosis was associated with a hematocrit-independent increase in blood viscosity characterized by a high plasma viscosity and normal or low hematocrit, conditions which were present both before primary surgery as well as at the time of the deep vein thrombosis diagnosis. Finally, in ovarian carcinoma patients prior to surgery, RBC aggregation, plasma viscosity, and platelet and fibrinogen concentrations were much higher than in control groups, while hemoglobin and hematocrit levels were lower in cancer patients than in healthy woman. Platelet, leukocyte, and fibrinogen concentrations were found to be significantly correlated with disease state, whereas plasma viscosity, RBC aggregation, hemoglobin and hematocrit were not. Patients who later developed deep vein thrombosis already had increased plasma viscosity prior to surgery. These findings confirm the presence of a hematocrit- and stage-independent hyperviscosity syndrome in untreated ovarian carcinoma patients.

<u>Drag-reducing agents</u>: Applied fluid mechanics describes how the addition of surfactants, long-chain polymers, small suspended solid particles, longish filaments, and micro bubbles reduces the abrasive action of turbulent flow in conveyor tubes (Weber et al.57 (1), 1991; Weber et al., 57(4), 1991) by converting turbulent into laminar flow (Fairchild et al., 1995). Medical research has shown that DRP may prove effective against atherosclerosis (Faruqui et al., 1987). It has also been suggested that the systemic administration of DRP may reduce the extravasation and metastasis of tumor cells (Kameneva, 2014). Resuscitation experiments on rats previously subjected to lethal hemorrhagic-shock revealed that injection of synthetic or natural, aloe-derived DRP, had substantial benefits due to an increase in near-wall hematocrit, or reduced plasma skimming (Macias et al., 2004). Concerning the negative impact of plasma skimming (increased near-wall velocity profile and hence wall shear stress), it was hypothesized that plasma skimming may cause a shear dependent release of vasodilators (Kameneva et al., 2004). It was argued (Marhefka, 2009) that the Toms effect (Toms, 1948), i.e., the reduction of hydrodynamic resistance in turbulent flow by DRP, would not be applicable in blood flow due to the low Reynolds numbers between 0.1 and 100 that characterize the circulatory system, well below the onset of turbulent flow. However, the authors had already demonstrated that the systemic administration of DRP at nanomolar concentrations reduces or eliminates the near-wall cell-free layer and increases blood flow in micro vessels. The DRP-induced occupation of near-wall space by RBCs (thereby increasing near-wall shear rates) may inhibit transendothelial tumor cell migration.

In areas near the internal blood vessel wall that have low shear stress, artherosclerosis and atherosclerosis may develop. Other studies (Sawchuk et al., 1999) confirmed that DRP are apt to increase wall shear stress, thereby preventing plaque formation through this mechanism.

Earlier work has also shown (Sevick et al., 1989, l.c.) that flow resistance increases with increasing tumor size, a finding which was attributed to a diminished Fahrsus effect (reduction in hematocrit in small vessels), a reduced Fahrffius-Lindqvist effect (reduction in blood viscosity in small vessels), and low shear rates in tumor vessels, which facilitate RBC rouleaux formation.

Furthermore, the administration of DRP was proposed as a novel hydrodynamic approach to treat coronary artery disease (Pacella et al., 2006).

<u>Polymer Conjugates:</u> Polymers have been used as therapeutic agents since the 1960s (Shelanski et al. 2006; Ravin et al., 1952) primarily as blood plasma expanders, wound dressings, and injectable or implantable depots. Modeling of pharmacologically active polymers is still based on Ringsdorf's propositions (Ringsdorf, 1975). Recently an exhaustive overview (Larson et al., 2012) of the design and chemistry for synthesizing polymer-drug conjugates was presented, with special emphasis on their potential applications and current development status as anticancer agents.

The main reasons for the interest in polymer-drug conjugates are their better water solubility compared to pure drugs (small molecule therapeutics), the fact that they increase the therapeutic index of anticancer agents via an enhanced permeability and retention effect, and their ability to avoid filtration via the kidneys, resulting in higher blood circulation time.

Despite the obvious potential advantages of polymer-drug conjugates as anticancer agents, there are serious obstacles to their use. Importantly, they must meet regulatory requirements for identity and purity. For example, studies must examine the metabolic fate of such conjugates, something that is difficult to determine for multicomponent systems. Among the different polymers being investigated as backbones for polymer-drug conjugates, the most widely used are still polyethylene glycol (PEG) and N-(2-hydroxypropyl)methacrylamide (HPMA) (some of these conjugates have already gained FDA approval). Although the biocompatibility of PEG has been established clinically it remains the drawback of its non-biodegradability, i.e., the fact, that it is subjected to renal filtration. Otherwise, its resistance against an uptake by the reticuloendothelial system is a significant advantage for this hydrophilic, highly hydrated polymer.

Several HPMA-based anticancer polymer-drug conjugates have reached clinical trial stage, and some biodegradable polymer-drug conjugates against prostate, breast, ovarian, colorectal, and lung cancers are currently under clinical evaluation (Larson et al., l.c.).

Since the pH of tumor tissue is about one order of magnitude lower than that found in healthy tissue (pH 6.5 vs. pH 7.4 respectively), research is also under way to develop site-specific drug delivery based on this pH difference.

<u>Mechanical degradation of DRA:</u> DRP and DRS are susceptible to flow-induced mechanical degradation (FIMD) due to collisions with RBCs or more rigid particles. This is a serious potential drawback to their use in medical applications.

From the literature one can extract the following facts about the FIMD of DRP and DRS:

FIMD of DRP: Although drag reduction increases with the molecular weight and length of the DRP, its susceptibility to flow-induced degradation increases as well. The rate of FIMD (mainly mid-scission of the molecule) increases with molecular weight and length. In so-called poor solvents (low solvation number) turbulent FIMD is higher at lower Reynolds numbers, and lower at high Reynolds numbers. Shear stability increases with solvation number. For constant wall shear stress and pipe diameter FIMD is directly proportional to the molecular weight and length of the DRP. For constant wall shear stress and DRP concentration FIMD is inversely proportional to pipe diameter. For constant wall shear stress and pipe diameter FIMD is inversely proportional to DRP concentration. For constant DRP concentration and pipe diameter FIMD increases with wall shear stress.

<u>FIMD of DRS:</u> Turbulent drag reduction by surfactants may result in a reduction of the friction factor of 70 - 80 %. Although DRS also undergo FIMD at high shear stress, they seem to be able to self-repair within seconds. Above a certain critical concentration, DRS solutions form characteristic rod-like micelles out of single surfactant molecules. Above a critical level of wall shear stress the rod-like micelles form a so-called shear-induced state (SIS) in which the micelles arrange themselves into larger structures aligned to the direction of flow. The SIS also represents the limit of achievable drag reduction by surfactants. The loss of drag reduction above the critical wall shear stress level is reversible below the critical wall shear stress level.

The mechanical degradation of drag-reducing agents has to be considered as a serious drawback for their use in medical applications. Interestingly however, natural Aloe Vera-derived DRP has shown lower rates of mechanical degradation.

## Methods

### Blood flow pattern

First we will investigate the question under which hydrodynamic circumstances it seems to be worthwhile to change the blood flow pattern near the vessel wall, i.e., in direction toward the vessel center. Our hydrodynamic approach is based on the threefold analogy between heat, mass, and momentum transfer (Welty et al., 1976; Baehr and Stephan).

The molecular transfer equations of Newton's law for fluid momentum at low Reynolds numbers (Stokes flow), Fourier's law for heat, and Fick's law for mass are very similar, since they are all linear approximations of the transport of conserved quantities in a flow field. At higher Reynolds numbers, the analogy becomes less useful due to the non-linearity of the Navier-Stokes equation or, more fundamentally, the general momentum conservation equation, but the analogy between heat and mass transfer still remains good according to (2)

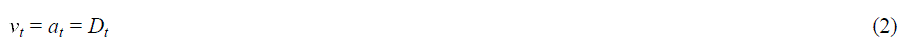

with
*v_t_* = turbulent kinematic viscosity; a_t_ = turbulent thermal diffusivity, and *D_t_* = turbulent mass diffusivity

The increase in mass transfer from laminar to turbulent flow is similar to the convective heat transfer. In the first case it is the ratio of the respective NuBelt numbers *Nu_t_/Nu_l_* which characterizes the difference while in heat transfer it is the ratio of the Sherwood numbers *Sh_t_/Sh_l_.* Eventually the effect of the momentum exchange on the shear stress can be characterized analogous by the corresponding products of the loss coefficient λ and the Reynolds number *Re*, i.e., *λ·Re* for tubes and ducts or *c_w_·Re* for flow around problems. Hence the increase in momentum exchange can be characterized by the ratio *λ·_t_·Re_t_ / λ_I_·Re_I_* with Re according to (3)

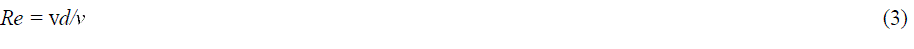

with v = flow velocity; *d* = tube diameter and *v =* kinematic viscosity

The Reynolds numbers change for the same tube when the flow pattern changes. If we consider first the laminar flow in a circular tube we get for the pipe friction factor (4)

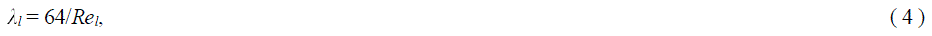

In fig. 2 we will represent graphically different flow pattern. This figure is a simplified adaptation of the well-known diagrams of Colebrook or Moody. A turbulent flow in a blood vessel with a smooth wall is shown at point 1. If by adding surfactants or DRP the flow pattern becomes laminar, flow resistance will be reduced. Assuming the same pressure drop and diameter, this results in a higher flow rate as well as an increased Reynolds number, as depicted at point 2.

**Fig. 2:**
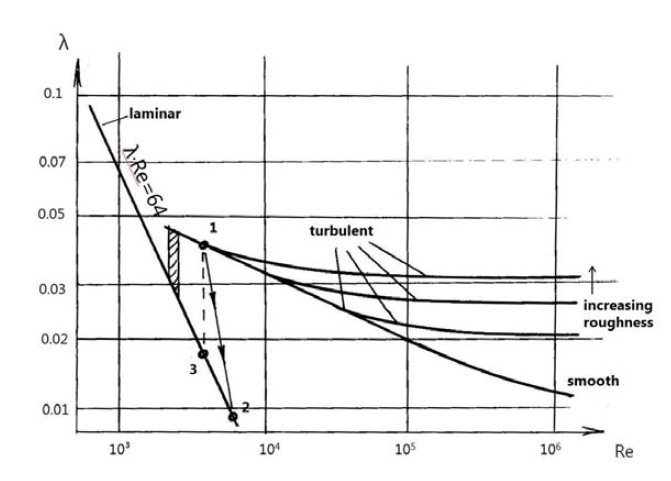
Shear stress vs. Reynolds number

The ratio *λ_2_Re_1_/λ_1_Re_2_* will then represent the achieved attenuation for the momentum and mass transfers respectively.

As can be seen from the straight line for laminar flow in the logarithmic plot of fig. 2, the value *λ_2_Re_2_* = 64 remains constant. Consequently, one can find a point 3 where *Re_3_* equals Re_1_; hence, with *Re_1_* = *Re_3_* the equation *λ_2_Re_2_ / λ_1_Re_1_= λ_3_Re_3_ / λ_1_Re_1_* is true. In this case obviously the Reynolds numbers cancel each other out and it remains the ratio *λ_3_* /*λ_1_* as the key flow parameter representing the achieved attenuation of momentum and mass transfer when surfactants or DRP are added. Henceforth we will refer to this ratio as the attenuation factor. The small hatched area represents a transition region where both states, laminar and turbulent flow, alternate.

One objection to this representation would be that it was developed for smooth and rough technical pipe walls. Blood vessels, however, are elastic and have ripples; hence the flow rate will depend on the axial coordinate and time. Nevertheless, these fluctuations can be considered as superpositions on the mainstream and will be of interest only for the determination of the true flow rate at each point in the vessel. For our purposes the interesting parameter is the determination of a mean flow rate or flow volume for a given pressure drop, since these values are necessary for the determination of the Reynolds number and hence of whether a flow is laminar or turbulent.

One can always find a mean diameter of a real blood vessel where for a given pressure drop the correct flow volume results. This vessel can further serve as sample vessel in order to find out to what degree a turbulent flow pattern can be affected. It is known (Ideltschik, 1975) that a weak waviness, i.e., large wavelength at small amplitude, doesn't have a decisive influence on the pressure drop. In the case of stronger ripples one can use a relative roughness.

In a healthy blood vessel the inner wall is smooth, pliant and mucous. The texture of the wall plays a role only when the small elevations are higher than the thickness of the laminar sublayer. Otherwise one can consider the surface as hydraulically smooth and this will be the case in real blood vessels.

Inserting in the numerator of our attenuation factor the Blasius formula (5)

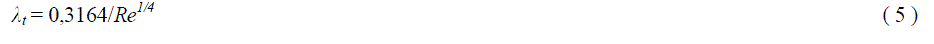

we get (6) as a measure to compare the turbulent and laminar mass and momentum transfers.

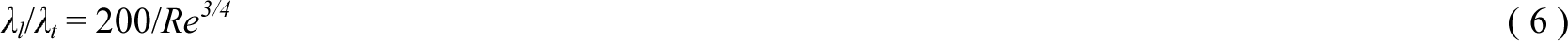

Yielding from (6) a value of 1/2 means halving the momentum and mass transfer. The same would be true for the heat transfer characterized by the Nusselt numbers which are available in some cases from literature. With a known Nusselt number one can derive finally also a mass transfer coefficient β defining a Sherwood number

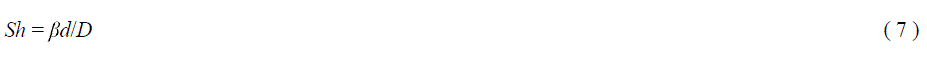

with *d* = tube diameter; *D* = diffusivity or diffusion coefficient; *Sh* = Sherwood number (also called the mass transfer Nusselt number, representing the ratio of convective to diffusive mass transport), and *β* = mass transfer coefficient

Although the Poiseuille law describes the laminar flow of an incompressible Newtonian fluid in a rigid pipe, it is often used as a first approximation to describe blood circulation. Most blood flow measurements are conducted in rotational viscosimeters (rheometers) of the Couette-or Searle-type, i.e., with fixed outer and rotating inner cylinders. Viscosities measured in rheometers are always mean or apparent viscosities while the true viscosity-gap distance relationship remains unknown. The term apparent viscosity is used in biomechanical literature when a linear falling shear rate in the gap between the both cylinders is assumed. It seems therefore to be worthwhile to describe at least qualitatively the flow conditions in a Searle-type rheometer (cf. fig. 3).

Studies indicate (Boron et al., 2005) that RBCs tend to accumulate in the center of blood vessels, leaving the wall region depleted (plasma skimming). This produces non-linear viscosity and hence shears rate profiles in the gap between the both cylinders. On the other hand the shear stress in the fluid is responsible for the transmission of the torque. Since the involved areas are outwardly bigger, the shear stress *τ(r)* falls in radius direction, cf. curve *τ(r)*. For a Newtonian fluid and laminar flow the shear rate would also be slightly sloping. For the non-Newtonian fluid blood and in the lack of true local viscosity data, i.e., assuming a constant apparent viscosity **η*_app_* (cf. dashed line), the derived shear rate function 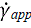 will also fall slightly from the inside to the outside of the gap (cf. curve 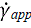). Due to the previously mentioned increased viscosity in the center, the true velocity v(r) (cf. curve v(r)), will show an inflection profile similar in shape to water in turbulent flow. For the same reason the increase in the velocity profile, i.e., the true shear rate 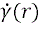, will be lower in the middle as opposed to the outward areas (cf. curve 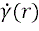).

**Fig. 3:**
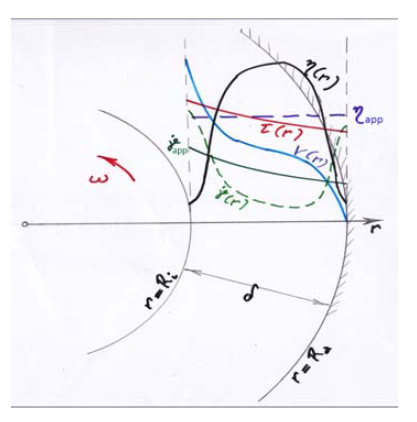
Couette flow with *r* = distance from the center *R_i_* and *R_o_* = pipe radius of the inner and the outer cylinders resp. *γ_app_* and *γ* (r) = apparent and true shear rates resp. *ω* = angular velocity **η*_app_* and **η*(r)* = apparent and true viscosities resp. *τ(r)* = true shear stress δ(r) = true velocity *δ* = gap width

In blood vessels the near-wall viscosity corresponds to the plasma viscosity of 1.2 cP, while in the vessel center we see values of about 10 cP due to the presence of RBC rouleaux (cf. curve **η*(r)* as noted above. Apart from RBC rouleaux formation, single RBCs also exhibit a certain resistance to shear through rotation about their centers of gravity, a process called tank treading (Boron et al., l.c.), which results in a higher dynamic viscosity level in the vessel center. Since plasma occupies about half of its volume, the frequently cited average viscosity of blood of 4.5 cP (Oertel, 2008 l.c.) is, therefore, well accounted for by hydrodynamics. This description of the curves is, of course, qualitative. Nevertheless, the general course of the different curves was drawn based on the results of bubble flow experiments and by taking into account the impact on viscosity of the local volume portion of the dispersed phase (Einstein, 1906 and 1911; Lilienblum, 1986).

### Lockhart-Martinelli procedure

In the second part of our hydrodynamic approach we argue for the application of the Lockhart/Martinelli (LM) procedure (Lockhart and Martinelli, 19491) to determine the pressure drop *Δp*_v_ in blood vessels. Our objective is to derive another mass transfer coefficient (analogous to the one described above) which characterizes the mass transfer between the center and the wall of blood vessels for different two-phase blood flow regimes. The different flow regimes are designated as “*ll*” when considering laminar flow patterns for both phases, “*tt*” for turbulent flow in both phases, and “*lt*” (or *“tl”*) for mixed laminar/turbulent flow conditions in the two phases. Aided by this new mass transfer coefficient, we will characterize the nutrient supply for tumor cells in blood vessels under different flow conditions, with special emphasis on the effect of DRA.

The starting-point for the LM procedure is Kirchhoff's law of the constancy of pressure drop in parallel branches. As previously discussed, in vessel constrictions (for example, those due to developed tumor cells at the wall) flow capacity is decreased, which in turn changes the wall shear rate, Re number and friction factor. Thus tumor cell obstacles may be said to cause a new representative viscosity as will be shown later on in 1table 2.

**Table 1:**
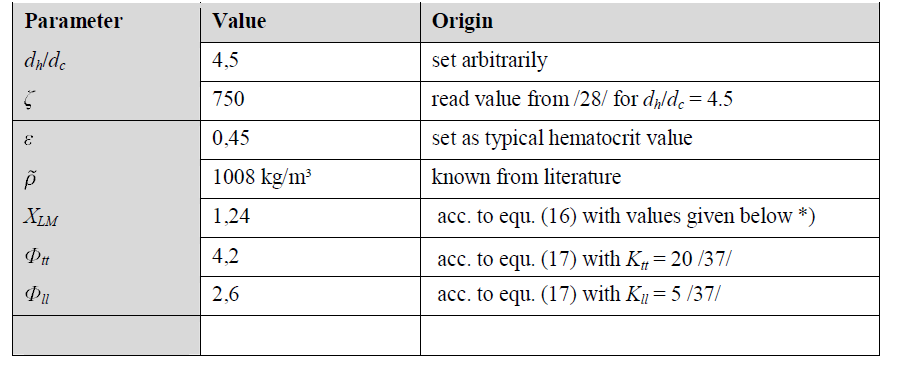
Calculations according to Lockhart-Martinelli

**Table 2:** Calculated flow parameters for healthy (h) and cancer infected (c) blood vessels

| Parameter | Dimension | Aorta | Arteries | Arterioles | Origin |
| --- | --- | --- | --- | --- | --- |
| $d$ | cm | 2 | 0.5 | 0.15 | set as typical values |
| $v_{\square_h}$ | m/s | 0.6 | 0.3 | 0.15 | acc. to (8) |
| $\dot{\gamma}_h$ | $s^{-1}$ | 240 | 480 | 800 | acc. to (14) |
| $\eta_{rep,h}$ | cP | 3.5 | 3.5 | 3.5 | read from the graph (fig. 4) |
| $Re_h$ | - | 3400<br>turbulent | 430<br>transition<br>turbulent/<br>laminar | 65<br>laminar | acc. to (1) |
| $\lambda_h$ | - | 0.04 | 0.05 | 0.34 | acc. to (4) for laminar flow or acc. to (5) for turbulent flow |
| $\tau_{w,h}$ | Pa | 0.84 | 1.7 | 2.8 | acc. to (12) |
| $v_{\square_c}$ | m/s | 0.014 | 0.017 | 0.031 | acc. to (19) |
| $l$ | m | 0.5 | 0.4 | 0.2 | set as typical values |
| $\dot{\gamma}_c$ | $s^{-1}$ | 5.6 | 25.5 | 165 | acc. to (14) |
| $\eta_{rep,c}$ | cP | 7.5 | 5.5 | 4.5 | read from the graph (fig. 4) |
| $\Phi$ | - | 4.2 (tt) | 3.4 (lt) | 3.0 (lt-II) | acc. to (17) |
| $\lambda_t / \lambda_l$ | - | 2.6 (tt) | 1.7 (lt) | 1.33 (lt – II) | acc. to (20) |

## Results and Discussion

Our hydrodynamic approach to cancer is based on the following reasoning:

Tumor cells need more nutrient-uptake than normal cells

Developed cancer cells within a tumor microvasculature may represent flow obstacles causing the formation of vortices or eddies, i.e., local turbulences.

Vessel bifurcation and bending as well as diameter reduction also produce local vortices with increased mass transfer rates and are therefore considered as potential docking points for circulating cancer cells.

In turbulent flow the mass transfer and hence the food supply are enhanced versus laminar flow.

The extra nutrient supply available in areas of turbulent flow is conducive to tumor cell growth.

Drag reducing agents can smooth down local vortices, thereby depriving tumor cells of their extra nutrient supply.

Drag reduction may possibly inhibit tumor growth or even lead to their starvation-induced death (Raffaghello et al., 2008) and, most importantly, it may reduce the risk of metastasis.

In order to be able to calculate the wall shear stress *τ_w_* using the experimentally-derived volumetric flow rate *V̇*, we first calculate the average flow velocity 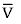 using (8)

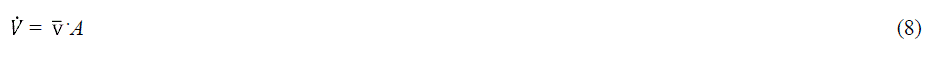

Where *A* = cross section area.

It should be noted that there is a great deal of literature concerning the measurement of blood viscosity. However, as previously mentioned, the rotational viscometers (rheometers) commonly used in blood rheology measure only apparent viscosity **η*_app_* since a lower plasma viscosity **η*_Pla_* exists at the vessel wall while at the center one finds a higher two-phase flow viscosity. Computer simulations take into account these differences in order to calculate the velocity profiles for laminar and turbulent flows respectively.

In a Hagen-Poiseuille flow one can replace computer simulations with simpler estimations (with errors in the order of 10 - 12 %, according to (/33/) under the assumption that blood is a Newtonian fluid. Using this method the experimentally derived apparent viscosity **η*_app_* can be used to determine the probability of RBC formation - an important calculation because RBC rouleaux closely resemble a typical turbulent flow pattern. Chandran /l.c./ attributes the increase in apparent viscosity **η*_app_* at low flow rates in the Whitmore diagram /34/ to the formation of RBC rouleaux.

Next we will compare the Hagen-Poiseuille flow to the Couette flow commonly employed in blood rheology. In order to extract the apparent viscosity **η*_app_* data from the Whitmore plot (fig. 4), it is necessary to calculate the apparent shear rate 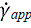. Note that while Whitmore uses the symbol 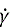 for the apparent rate of shear, we designate it in the following as 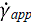.

The comparison of two non-Newtonian flows using the approximate laminar velocity profiles of a Newtonian fluid is a specifically hydrodynamic type of analysis, something which was previously suggested as a possible procedure by Chandran et al. (2007).

The Casson equation depicted also in fig. 4 can be derived from the general form 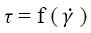 considering as well the yield stress *τ*_o_ as the superposition of a nonlinear 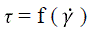 dependence with increasing shear rate. This can be represented by a square root function, which is superimposed on a linear function according to (9)

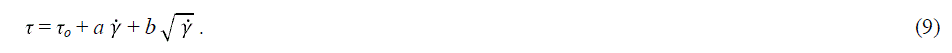

where *τ* = shear stress, *τ_o_* = yield stress, *a* and *b* = constants, and 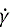 = shear rate.

**Fig. 4:**
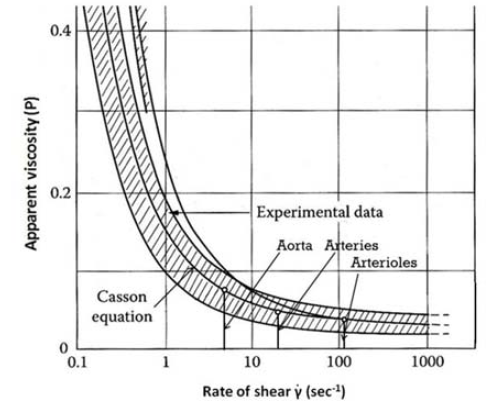
Plot of apparent viscosity as function of shear rate (extract redrawn from Chandran et al. (2007) with permission)

Another form of the Casson equation (derived in /5/) combines shear stress *τ*, plasma viscosity *λp*, = 1.2 cP and shear rate γ where best fit with experimental results is achieved according to (10)

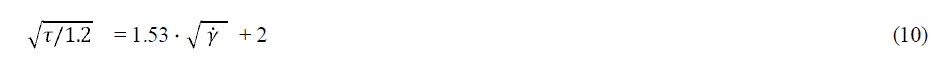

Chandran et al. (2007) compares furthermore the Casson equation with the flow laws of Newton and Bingham, and with the Power law (cf. fig. 5).

**Fig. 5:**
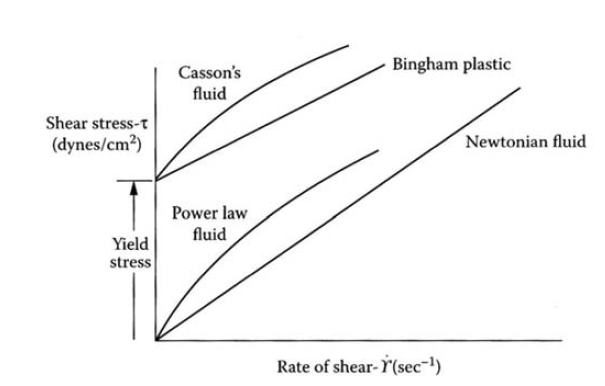
Shear stress vs. shear rate plots for Newtonian and non-Newtonian fluids

In our calculations going forward we will use the apparent shear rate *f* obtained from rheometer measurements, keeping in mind that the true dynamic viscosity is lower near the plasma dominated wall and higher in the vessel center due to RBC accumulation.

Fig. 6 illustrates the Hagen-Poiseuille flow as a model for blood vessel circulation, in which we depict in a qualitative manner how key flow parameters (wall shear stress τ_*w*_, apparent viscosity **η*_app_*, and apparent shear rate 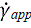) depend on the distance from the wall *r*.

**Fig. 6:**
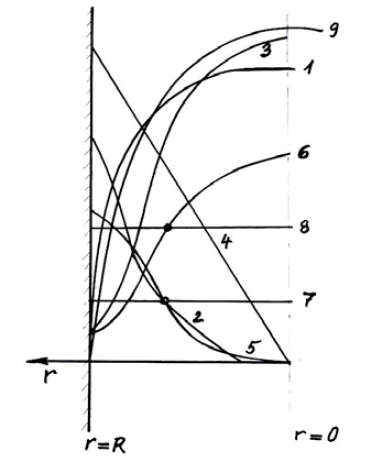
Qualitative dependence of key flow parameters on the distance from the wall in Hagen-Poiseuille pipe flow 1 true velocity 2 true yD (r) with yD = shear rate 3 true viscosity 4 shear stress 5 apparent yD (r) 6 apparent viscosity 7 representative yD (r) 8 representative viscosity 9 velocity calculated Newtonian

As one can see from fig. 6 the shear stress *τ* decreases linearly from the wall towards the vessel center. The apparent viscosity **η*_app_* in turn increases in a non-linear fashion in the same direction. When the exact **η*_app_* (*r*) dependence is unknown, we use a representative viscosity **η*_rep_.* The same approximation is used for the true (unknown) shear rate 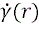 dependence, i.e. we use a representative shear rate *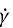_rep_*.

To perform the calculation as outlined, it is necessary to find a representative viscosity to which a representative shear rate can be attributed, as recommended in /3 5/. The resulting velocity profile is then depicted qualitatively by curve 1.

If we consider in a Hagen-Poiseuille pipe flow a certain coaxial cylindrical volume element at a given time we can equate the pressure drop per cross section area *Δp_v_* to the wall shear stress τ_w_. The pressure drop results than from the friction factor *λ*. combined with the mean velocity 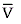 according to (11) with *l* = tube length

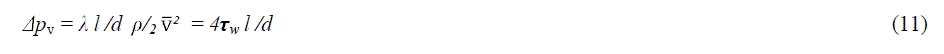

With *λ*. = 64/*Re* from (4) one gets (12) when specifying an initially unknown representative viscosity of the blood

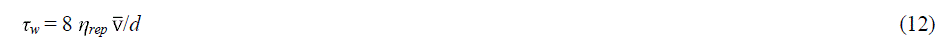

with **η*_rep_* = representative viscosity as depicted in fig. 6.

Comparing (11) with (12) from Couette flow according to (13)

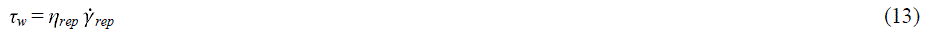

with *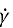_rep_* = representative shear rate
one gets for Hagen-Poiseuille flow at the inner wall (14)

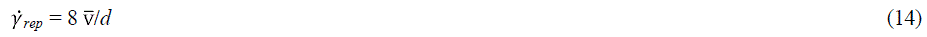

and in turn from the Whitmore diagram the value **η*_rep_* as wanted

Next one has to consider that in Hagen-Poiseuille flow the shear stress τ is not constant but decreases towards the center and the same is true for 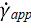,. In recent literature (Chandran et al., 2007) equation (15) is recommended as basis for further calculations.

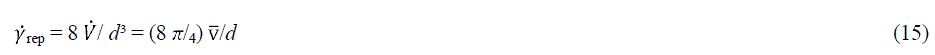

Following this procedure we calculated first acc. to (15) the for the cross section representative shear rate *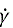_rep_*data of Table 2 and read then with help of these *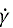_rep_* data from the Whitmore diagram (fig. 4) the corresponding apparent viscosity data *rj_rep_* given in Table 2.

Concerning the first aim, i.e., to elucidate under which circumstances a change in the blood flow pattern seems worthwhile the data in tables 1 and 2 can be interpreted as follows (the index *h* stands for healthy, and the index *c* for cancer cell infected vessels resp.):

1. Developed tumor cells within a tumor microvasculature or attached to the inner vessel wall represent flow constrictions giving rise to decreased mean blood flow velocity, decreased wall shear rate, and decreased Re numbers, however an increased friction factor.
2. Despite the considerable decreased Re numbers in tumor infected vessels (Re_c_ << Re_h_) the decreased blood flow favors RBC aggregation, i.e., formation of local vortices with enhanced mass transfer towards the wall. This enhanced mass transfer assists in creating an extra-food supply to tumor cells thus turning the food-uptake situation for tumor cells in a fatal vicious circle.
3. Injection of DRA may inverse the flow situation in shifting the flow pattern in direction of a healthy blood circulation system.

Next we will develop and discuss the arguments for the second aim, i.e., the applicability of the Lockhart/Martinelli (LM) procedure in order to determine the pressure drop *Δp_v_* in blood vessels from the Lockhart-Martinelli-diagram (fig. 7).

**Fig. 7:**
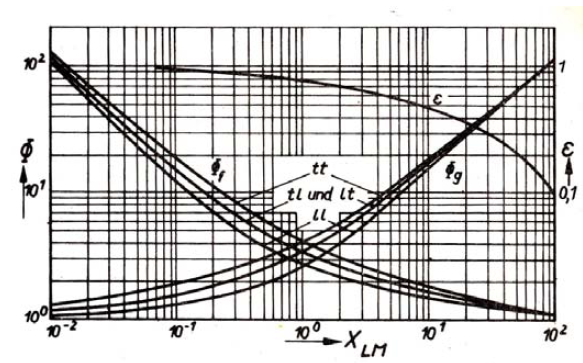
Lockhart-Martinelli diagram

For the calculation of the LM parameter it is important to choose the proper viscosity values acc. to (12). While whole blood has a dynamic viscosity of 4.5 cP, we have chosen for the plasma dominated near wall region a viscosity of 1,2 cP, and assumed for the inner vessel region due to the rouleaux formation a value of 10 cP.

As already pointed out at low shear rates the primary sporadic scattered rouleaux may develop ramifications called secondary rouleaux (Kulicke, 1986, l.c.). This explains together with the fibrinogen adhered to the wall and to the rouleaux fibrinogen filaments the increased viscosity at low shear rates and also the yield shear stress. Thus arises a slip between plasma and the rouleaux-fibrinogen cluster which causes strong local vortices even in the absence of fully developed turbulent flow. These cluster-induced eddies near the wall have to be attributed as localized “tt” domains. However, at Re numbers around 100 and below according to LM blood flow will be treated as ‘***It***’. Hence, one has to keep in mind that mass transfer under these circumstances is governed by local flow perturbations although a first look on the rather low Re numbers seems to point on laminar flow conditions. Obviously only a closer look at local flow pattern solves the apparent contradiction.

Increasing shear rates dissolve the rouleaux and the RBCs occupy the entire vessel cross-section. However, depending on vessel diameter the well-known phenomena of tank treading, plasma skimming, and deformation of RBCs to bullets may occur (Boron et al., 2005, l.c.). The flow condition for tank treading will be discussed in terms of “***ll***” (i.e., flow is constrained laminar although Re numbers point on turbulence).

With the designations pla for plasma and rbc for RBCs we define the Lockhart-Martinelli parameter X_LM_ according to (16) (Köhler, 1996).

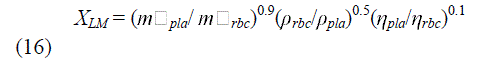

with

*m□_pla_* and *m□_rbc_* = mass flow rates of plasma and RBCs resp.

*ρ_pla_* and *ρ_rbc_* = densities of plasma and RBCs resp., and

**η*_pla_* and **η*_rbc_* = viscosities of plasma and RBCs resp.

For the densities of erythrocytes *ρ_rbc_* and plasma *ρ_pla_* we have chosen 1100 g/l and 1028 g/l respectively.

Provided the slippage between plasma and RBCs is small the ratio of the mass flow rates *m□_pla_/m□_rbc_* equals about the corresponding masses ratio *m_la_/ m_rbc_.* This masses ratio in turn is due the similar densities of the plasma and the RBC phase almost equal to the ratio of the volumes of the both phases, i.e., the hematocrit. Hence, one can use the hematocrit ε = 0.45 as a first approximation for the ratio of the mass flow rates. However, one should point out that in arterioles where the vessel diameter is of the same order as the RBC dimension may occur a slip between both phases which can even become a source of local eddies.

Next, X_LM_ will be corrected according to (17) (Chrisholm, 1967)

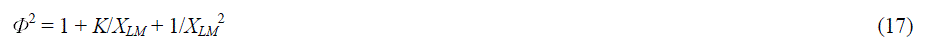

with
*K_ll_* = 5 and *K_tt_* = 20 (*K* = Chisholm correction factor)
in order to arrive at a corrected pressure drop *Δp_v_* (18)

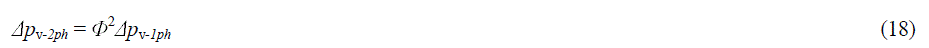

The pressure loss due to a decreased (tumor cell constricted) vessel diameter can be characterized by the pressure drop coefficient zeta (ζ (Ideltschik, 1975, l.c.). With help of (18) one calculates pressure drops *Δp_v-h_* and *Δp_v-c_* for healthy (index *h*) and cancer infected (index *c*) vessels. The reduced (by diameter constriction) blood flow velocity is calculated according to (19) under consideration of the discussed above two-phase flow pattern (“*tt*” for turbulent rouleaux-burdened and “*ll* for DRP-laminarized flow pattern)

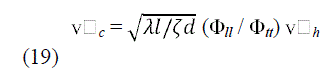

In front of the constriction one has a blunted flow which exhibits despite a decreased mean velocity a strongly enhanced mass transfer in local vortices. DRP injection should not only reduce the friction factor λ. but also the pressure drop coefficient zeta (ζ)

Applying the depicted procedure we calculated with help of known hemodynamic data from literature the respective v□_*c*_ and v□_*h*_ data depicted in table 2. We used an arbitrary constrictions ratio of *d_h_*: *d_c_* = 4.5 : 1 resulting at the given Re number in a *ζ* value of 750 (Ideltschik, 1975) at a hematocrit of 45% and we have chosen arbitrary but reasonable diameter (*d*) and velocity values (v□_*h*_ and v□_*c*_) for the three vessel types aorta, arteries and arterioles (cf. table 2).

The results can be interpreted as follows: Especially in big vessels addition of RDA may result in considerable decrease of the ratio of the turbulent and laminar friction factors λ_*t*_/λ_*l*_ (20)

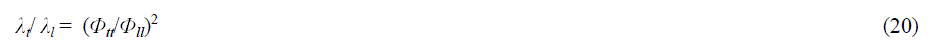

It is therefore worthwhile to prevent RBC aggregation in order to reduce enhanced mass transfer.

Fig. 8 depicts in a qualitative manner the assumed velocity profiles of blood flow with and without addition of surfactants and DRP respectively. The three curves can be interpreted as follows: Without the addition of DRA at low velocities (e.g., at 0.02 m/s or less) the blood flow in the vessel center resembles a RBC-plug with blunted profile. Adding surfactants generates better RBC mobility, which at best results in a homogeneous flow pattern in the center. Near the wall, however, the profile is steeper due to the predominance of plasma, although it is less steep than with RBC rouleaux. The higher velocity in the vessel center leads automatically to a lower velocity near the wall. The addition of DRP produces principally the same effect as surfactants. Near the wall, however, viscosity will be enhanced due to the alignment of DRP filaments in the direction of blood flow. The velocity profile is less pronounced when compared to the situation without additives or to the profile resulting from the addition of surfactants, which automatically generates a higher velocity in the center.

**Fig. 8:**
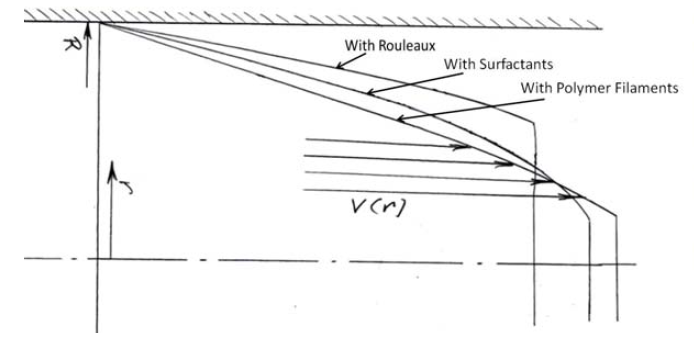
Velocity profiles

## Conclusion and critics

One major limitation of this paper relates to the compromises that were necessary in order to apply the principles of classical hydrodynamics to the analysis of flow in the body's circulatory system. Unlike the rigid tubes for which these principles were developed, blood vessels are soft, flexible, extensible, deformable, and able to transmit undulations caused by the heart's pumping action.

Despite the considerable progress that has been made using computer simulations based on viscoelastic models of blood vessels (Sequeira et al., 2014; Robertson et al., 2008) the complexity of blood rheology still prevents a complete description of circulation. For example, vessel walls show complex stress-strain characteristics. Their composite structure, consisting of elastin and collagen, makes the walls elastic; however, their elasticity strongly depends on applied stress - their stiffness increases with stress. Even if the plasma is replaced by an isotonic solution the suspension behaves in a non-Newtonian fashion that is attributable to the RBC flexibility. Vessel size and geometry influence flow resistance. While bends alter the flow pattern in large vessels, junctions cause hematocrit changes in small vessels due to plasma skimming. However, in large vessels junctions give also rise to localized turbulence. Our novel approach - to apply the Lockhart/Martinelli procedure to the two-phase blood flow - seems therefore to be warranted owing to its simplicity.

The proposal to inject drag-reducing agents into cancer-infected blood vessels in order to prevent the docking of circulating cancer cells and possibly even control cancer cell growth is based on classical hydrodynamics theory.

Previous work on DRA injection was focused on improving blood supply in areas where it was impaired. Our proposed hydrodynamic approach, on the other hand, is aimed at using drag-reducing agents to reduce mass transfer in order to withhold a surplus nutrient supply to tumor cells.

Although initial studies of the mechanical degradation of DRP have taken place (Marhefka and Velankar et al., 2008) additional data on their blood circulation time and possible unwanted side effects are necessary.

Our proposal to prevent tumor metastasis by laminarising local turbulent blood flow patterns implies a novel type of cancer aftercare that requires verification using animal models before advancing to clinical trials involving human patients.

## Acknowledgements

The authors want to thank K. Pinkert and D. Butler for valuable help.

## List of abbreviations in tables 1 and 2 resp.

*d_h_/d_c_*: = ratio of diameters in healthy (h) and cancer infected (c) vessels,
*ζ*: =pressure drop coefficient acc. to /28/
*ε*: =hematocrit
*ρ͂*: =mean blood density
*X_LM_*: = Lockhart-Martinell (LM) factor
*Φ_tt_*: = corrected LM factor for turbulent flow in both phases
*Φ_ll_*: = corrected LM factor for laminar flow in both phases
*d*: = diameter of blood vessels
v*□_h_*: = mean blood flow velocity in healthy vessels
*γ̇_h_*: = shear rate in healthy vessels
**η*_rep_,h*: = representative viscosity in healthy vessels
*Re_h_*: = Reynolds number in healthy vessels
*λ_h_*: = friction factor in healthy vessels
*τ_w,h_*: = wall shear stress in healthy vessels
*v□_c_*: = mean blood flow velocity in cancer infected vessels
*l*: = tube length
*γ̇_c_*: = shear rate in cancer infected vessels
**η*_rep,c_*: = representative viscosity in cancer infected vessels
*Φ*: = corrected acc. to (17) Lockhart-Martinelli factor
*λ_t_* / *λ_l_*: = attenuation factor = ratio of friction factors for turbulent (t) and laminar (l) flows resp.

